# WDR5 stabilizes actin architecture to promote multiciliated cell formation

**DOI:** 10.1101/153361

**Authors:** Saurabh S. Kulkarni, John N. Griffin, Karel F. Liem, Mustafa K. Khokha, on behalf of the PCGC Investigators

## Abstract

**HIGHLIGHTS:**

- WDR5 has an H3K4 independent role in the formation of multiciliated cells.
- WDR5 controls apical cell expansion, basal body patterning, and ciliogenesis in multiciliated cells.
- WDR5 localizes near the ciliary base where it connects basal bodies to F-actin.
- WDR5 stabilizes the apical actin network in multiciliated cells.

**SUMMARY:** The actin cytoskeleton is critical to shape cells and pattern intracellular organelles to drive tissue morphogenesis. In multiciliated cells (MCCs), apical actin forms a lattice that drives expansion of the cell surface necessary to host hundreds of cilia. The actin lattice also uniformly distributes basal bodies across this surface. This apical actin network is dynamically remodeled, but the molecules that regulate its architecture remain poorly understood. We identify the chromatin modifier, WDR5, as a regulator of apical F-actin in multiciliated cells. Unexpectedly, WDR5 functions independently of chromatin modification in MCCs. Instead, we discover a scaffolding role for WDR5 between the basal body and F-actin. Specifically, WDR5 binds to basal bodies and migrates apically, where F-actin organizes around WDR5. Using a monomer trap for G-actin, we show that WDR5 stabilizes F-actin to maintain apical lattice architecture. In summary, we identify a novel, non-chromatin role for WDR5 in stabilizing F-actin in multiciliated cells.

**IN BRIEF:** Kulkarni et al discover a chromatin independent function for WDR5 in multiciliated cell formation. WDR5 localizes to the base of cilia and functions as a scaffold between the basal bodies and the apical actin lattice. There, WDR5 stabilizes the actin lattice that allows multiciliated cells to expand their apical surface, pattern basal bodies, and generate hundreds of cilia.

## INTRODUCTION

Cilia play a central role in development and disease as they perform multiple sensory, motility and signaling functions (Oh and Katsanis, 2012; Bettencourt-Dias, et al., 2011; Sharma, et al., 2008). Cilia in multiciliated cells (MCCs) are specifically important for creating mechanical force to drive extracellular fluid flow, which is critical for clearing mucus in the airways, for transporting the egg through the oviduct, and for circulating cerebrospinal fluid in the cerebral ventricles (Oh and Katsanis, 2012; Bettencourt-Dias, et al., 2011; Houtmeyers, et al., 1999). Despite the importance of multiciliated cells (MCCs) in health, our understanding of ciliogenesis remains incomplete. While the formation of a single cilium is complex, in a MCC, this complexity is compounded to generate the hundreds of oriented cilia that beat in a coordinated fashion. First, a specialized structure called the deuterosome produces hundreds of centrioles that become the basal bodies necessary to seed many cilia (Klos Dehring, et al., 2013; Tang, 2013; Zhao, et al., 2013). Second, these basal bodies associate with vesicles that migrate and dock to the apical cell surface (Burke, et al., 2014; Vladar and Axelrod, 2008). Finally, ciliary axonemes protrude from the basal bodies and elongate to form functional cilia (Ishikawa and Marshall, 2011; Pedersen and Rosenbaum, 2008). The challenge for hundreds of cilia in MCCs is coordinating beating to effectively drive extracellular fluid flow. To do so, the cilia must be aligned within the epithelial plane, which is established by the rootlets and basal feet, which are attached to the basal body (Kunimoto, et al., 2012; Mitchell, et al., 2009).

Actin is a major driver of MCC formation (Sedzinski, et al., 2016; Werner, et al., 2011; Vladar and Axelrod, 2008). First, establishing a mucociliary epithelia such as in the *Xenopus* embryonic epidermis requires actin dependent radial intercalation of nascent MCCs (Walck-Shannon and Hardin, 2014). Specifically, nascent MCCs are first specified in deeper layers but migrate apically to insert in the surface (Sedzinski, et al., 2016; Werner, et al., 2014; Stubbs, et al., 2006). Then, once inserted, apical actin forms a lattice that generates the cell autonomous 2D force to expand the MCC’s apical surface, which is necessary to host hundreds of cilia (Sedzinski, et al., 2016). In addition, basal bodies are evenly distributed within this apical actin lattice across the cell surface (Antoniades, et al., 2014; Werner, et al., 2011). Finally, the planar polarization of cilia may require this apical actin lattice (Werner, et al., 2011; Park, et al., 2008). Therefore, F-actin plays a central role in the formation, patterning, and subsequent function of cilia in MCCs. However, our understanding of the molecular regulation of actin assembly in MCCs remains rudimentary (Sedzinski, et al., 2017; Sedzinski, et al., 2016). Specifically, how the actin lattice architecture is defined and maintained is not understood.

Here, we show that WDR5 is a key regulator of the apical organization of actin in MCCs. WDR5 is a core subunit of the human histone H3 Lys4 methyltransferase (H3K4MT) complexes that are essential for chromatin modification and transcriptional regulation (Trievel and Shilatifard, 2009; Patel, et al., 2008b; Dou, et al., 2006; Wysocka, et al., 2005). In particular, WDR5 is a highly conserved scaffolding protein essential for the association of RbBP5, ASH2L, and mDPY-30 with MLL1 (Odho, et al., 2010; Trievel and Shilatifard, 2009; Patel, et al., 2008b). The assembly of this complex is essential for H3K4MT activity (Dharmarajan, et al., 2012; Wysocka, et al., 2005). WDR5 acts as a scaffold between MLL1 and other members of the complex. WDR5 has an arginine-binding cavity that interacts with the arginine-containing WIN (WDR5- interacting) motif of MLL (Dharmarajan, et al., 2012; Patel, et al., 2008a). In fact, a mutation in the WIN motif of the MLL protein (R3765A) or the arginine-binding cavity in WDR5 (S91K) disrupts assembly and activity of the H3K4MT complex (Patel, et al., 2008b). Nearly all studies of WDR5 focus on its nuclear function regulating H3K4 methylation, and while WDR5 has been localized outside the nucleus, a cytoplasmic role is not well defined (Bailey, et al., 2015; Wang, et al., 2010).

Our studies began when WDR5 was identified from a genomic analysis of congenital heart disease (CHD) and heterotaxy patients (Zaidi, et al., 2013). Heterotaxy (Htx) is a disorder of left-right patterning that can have a severe effect on cardiac patterning and function (Sutherland and Ware, 2009). A patient with a *de novo* missense mutation (K7Q) in WDR5 exhibited a conotruncal defect and a right aortic arch (normally the arch is on the left) (Zaidi, et al., 2013). How this chromatin modifier might affect a specific phenotype such as cardiac development was unknown. Depletion of WDR5 in *Xenopus* recapitulates a left-right patterning defect with abnormal ciliogenesis in the left-right organizer (Kulkarni *et al* submitted).

Importantly, patients with CHD and Htx suffer from high postsurgical morbidity and mortality often associated with respiratory complications, a result of cilia dysfunction in MCCs (Garrod, et al., 2014; Nakhleh, et al., 2012; Swisher, et al., 2011; Kennedy, et al., 2007). Since a number of genes are essential for ciliogenesis in both monociliated cells (as in the left-right organizer) and MCCs (mucociliary epithelia), here, we examined if WDR5 also plays an important role in ciliogenesis in MCCs (Casey, et al., 2015; Nakhleh, et al., 2012; Becker-Heck, et al., 2011; Merveille, et al., 2011; Kennedy, et al., 2007; Tan, et al., 2007). Wdr5 is critical for ciliogenesis in the MCCs, but, unexpectedly, *independently* of chromatin modification. In fact, we discover that Wdr5 is localized near the base of cilia. Interestingly, Wdr5 regulates the enrichment of apical actin, and MCCs depleted of Wdr5 fail to expand their apical surface and lose planar distribution and organization of basal bodies. Using high-resolution confocal imaging, we show that Wdr5 localizes between F-actin and the basal bodies suggesting an alternative scaffolding role between these two structures. Further, using live imaging, we find that Wdr5 first associates with basal bodies deep in the cytoplasm and migrates apically. Upon reaching the apical surface, Wdr5 dynamically interacts with F-actin, where F-actin appears to organize around Wdr5. Using a monomer trap experiment, we show that Wdr5 is necessary to stabilize these actin polymers. Taken together, we discover a non-chromatin function for WDR5, where it stabilizes the actin lattice architecture to promote formation and function of MCCs.

## RESULTS

### WDR5 is essential for cilia in MCCs

In a separate report, we describe the role of WDR5 in left-right patterning via regulation of ciliogenesis in the left-right organizer (Kulkarni *et al* submitted). Because respiratory complications are an established co-morbidity in CHD patients with Htx (Li, et al., 2015; Nakhleh, et al., 2012; Swisher, et al., 2011), we also examined the cilia in the MCCs of the *Xenopus* embryonic epidermis, which is the focus of this study. In *Xenopus*, the embryonic epidermis has an array of MCCs that provide an ideal model to study cilia assembly and function (Werner and Mitchell, 2013). Wdr5 depletion resulted in a dramatic loss of cilia, specifically fewer and shorter cilia in MCCs (Figure 1A), suggesting that WDR5 may play an important role in mucociliary clearance. To this end, we examined cilia-driven extracellular fluid flow over the surface of the *Xenopus* embryo as a test of cilia function in MCCs. We visualized the flow by adding latex microspheres (beads) to the culture media followed by time-lapse imaging of bead movement. In WT embryos, cilia-generated flow was brisk (Figure 1 B,C, Video S1). Wdr5 depletion led to a complete loss of cilia generated fluid flow as expected for a dramatic loss of cilia (Figure 1B,C, Video S1).

**Figure 1:**
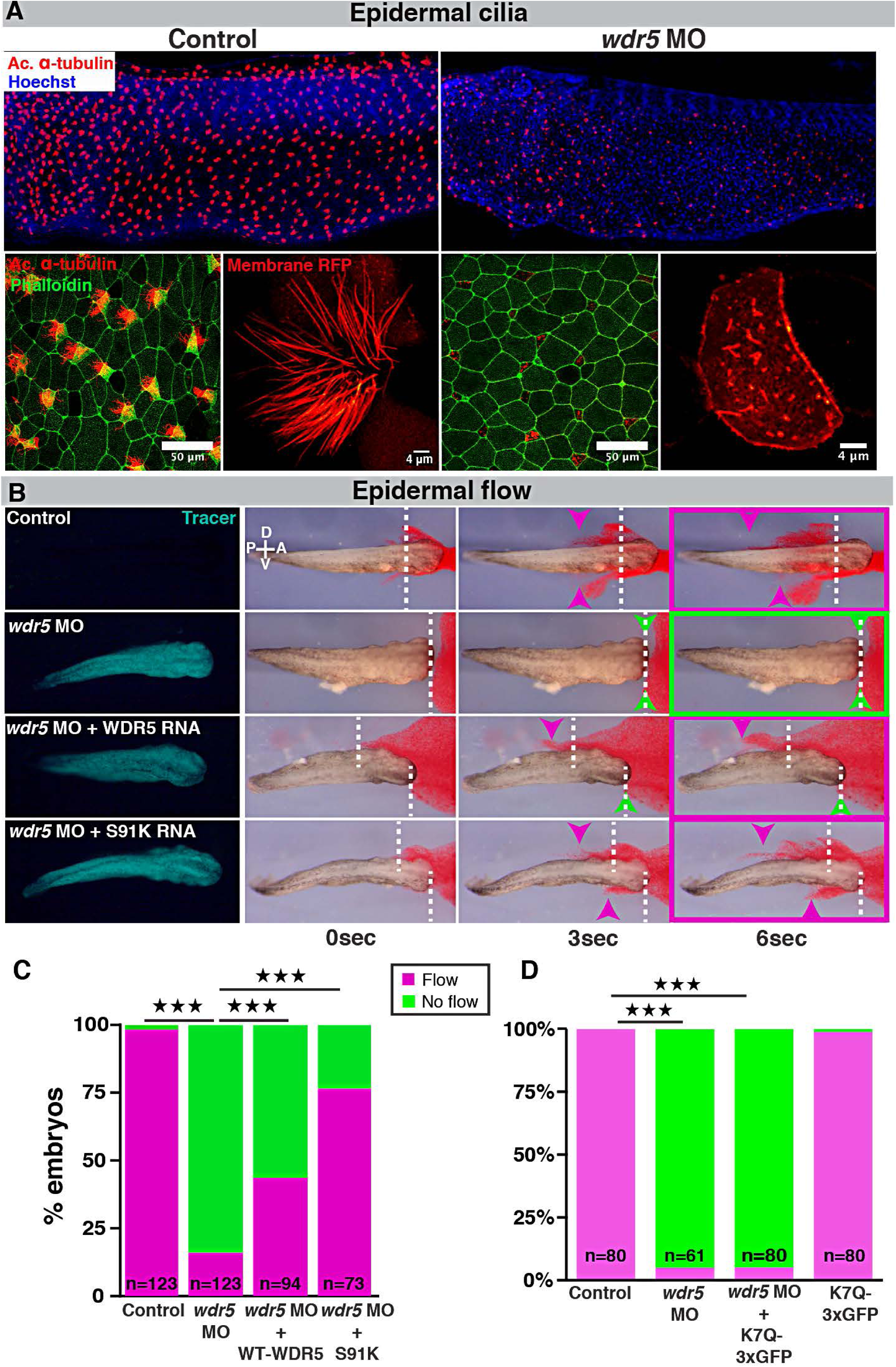
Wdr5 regulates ciliogenesis in MCCs independently of H3K4MT. (A) *X. tropicalis* epidermal MCCs marked either by anti-acetylated a-tubulin (red) or membrane-RFP, actin is labeled with phalloidin and nuclei with Hoechst. Uninjected control embryos on left and embryos injected at one cell stage with *wdr5* MO on right. (B) Cilia driven epidermal flow is visualized using red microbeads over the period of 6 seconds (see Movie S1). Magenta and green arrowheads indicate bead displacement from starting point (dashed lines). Green fluorescent protein traces the morpholino and RNA. Embryos were injected with *wdr5* MO, *wdr5* MO + human WT WDR5 RNA, *wdr5* MO + human mutant S91K WDR5 RNA and compared to uninjected controls. (C, D) Percentage of embryos with present or absent cilia driven epidermal flow. (c) Embryos were uninjected controls or injected with *wdr5* MO, *wdr5* MO + human WT WDR5 RNA, *wdr5* MO + human mutant S91K WDR5 RNA. (D) Embryos were uninjected controls or injected with *wdr5* MO or *wdr5* MO + human WDR5 (K7Q)-3xGFP RNA. n = number of embryos. ⋆⋆⋆ indicate statistical significance at *P* < 0.0005. See also Figure S1; Movies S1-S4.

To test the specificity and efficiency of our Wdr5 depletion, we employed multiple tests. First, we detected a reduction in Wdr5 protein in morphants by Western Blot that is partially rescued by injecting human wildtype (WT) WDR5 mRNA (Figure S1A). We tested the specificity of our Wdr5 antibody by serial dilution of a blocking peptide that reduced Wdr5 signal on the Western Blot (Figure S1B). Third, injecting a scrambled MO did not result in any change in cilia driven flow compared to WT embryos (Figure S1C). Fourth, co-injecting a wildtype human WDR5 mRNA in *wdr5* morphants rescued the flow defect, confirming the specificity of our knockdown (Figure 1B,C, Video S1).

The WDR5 patient had a *de novo* K7Q missense mutation (Zaidi, et al., 2013); however, whether this represented a loss of function allele for mucociliary clearance was unclear. To test this hypothesis, we attempted to rescue the loss of cilia-driven fluid flow phenotype in *wdr5* morphants with the K7Q WDR5 variant. In contrast to WT WDR5, the K7Q WDR5 variant failed to rescue ciliary flow defects in *wdr5* morphants supporting the hypothesis that it is a loss of function allele (Zhu, et al., 2017) (Figure 1D). In sum, we conclude that the Wdr5 depletion by MO is specific, and Wdr5 is essential for ciliogenesis in the MCCs.

### WDR5 has an H3K4 *independent* role in ciliogenesis in MCCs

WDR5 is a core subunit of the MLL/SET1 H3K4MT complex and has been studied extensively for its role in histone methylation, chromatin modification, and transcriptional regulation (Dharmarajan, et al., 2012; Odho, et al., 2010; Trievel and Shilatifard, 2009; Patel, et al., 2008a; Patel, et al., 2008b; Dou, et al., 2006; Gori, et al., 2006; Wysocka, et al., 2005). Therefore, we decided to examine if WDR5 regulates ciliogenesis in MCCs via H3K4 methylation.

Previous biochemical studies have identified that one face of the WDR5 beta-propeller binds to the WIN motif of the MLL/SET catalytic subunit (Dharmarajan, et al., 2012; Trievel and Shilatifard, 2009; Patel, et al., 2008a). A mutation in the arginine-binding cavity (S91K WDR5) disrupts the assembly and activity of the H3K4MT complex (Patel, et al., 2008b). Before embarking on RNA or ChIP-sequencing experiments to identify WDR5/ H3K4 targets, we tested the human S91K WDR5 variant for rescue in MCCs of *wdr5* morphants. We began by analyzing cilia generated extracellular fluid flow. To our surprise, co-injecting the S91K mutant RNA rescued the flow defect in *wdr5* morphants strongly suggesting that Wdr5 may be playing an H3K4 independent role in cilia (Figure 1B,C, Video S1).

### WDR5 is localized near the base of the cilium

To identify an alternative, non-H3K4 role for Wdr5 in MCCs, we overexpressed a human WDR5- GFP construct in *Xenopus* embryos. As expected, we did detect GFP signal in the nucleus but also in a punctate pattern near the apical surface of the MCCs, reminiscent of basal bodies (Figure 2A,B). We verified that WDR5-GFP is functional and localized appropriately by injecting WDR5-GFP mRNA in *wdr5* morphants to rescue cilia driven flow defects (Figure S1D). Even then, wary of overexpression artifacts of GFP-tagged protein, we sought to confirm this Wdr5 localization near the base of cilia by immunofluorescence. Using an anti γ-tubulin and an anti-WDR5 antibody, we found that WDR5 localizes near the base of cilia in the MCCs of the *Xenopus* epidermis (the membrane localization of the WDR-GFP may be non-specific) (Figure 2C). To see if this finding generalizes to mammals, we examined the MCCs of mouse tracheal epithelia. Again, we found that WDR5 is localized near the ciliary base in mammalian MCCs (Figure 2D), and the signal can be eliminated in the presence of a blocking peptide (ratio 1:2 antibody: blocking peptide) indicating specificity. Therefore, in *Xenopus* and mouse MCCs, WDR5 is located near the base of cilia.

**Figure 2:**
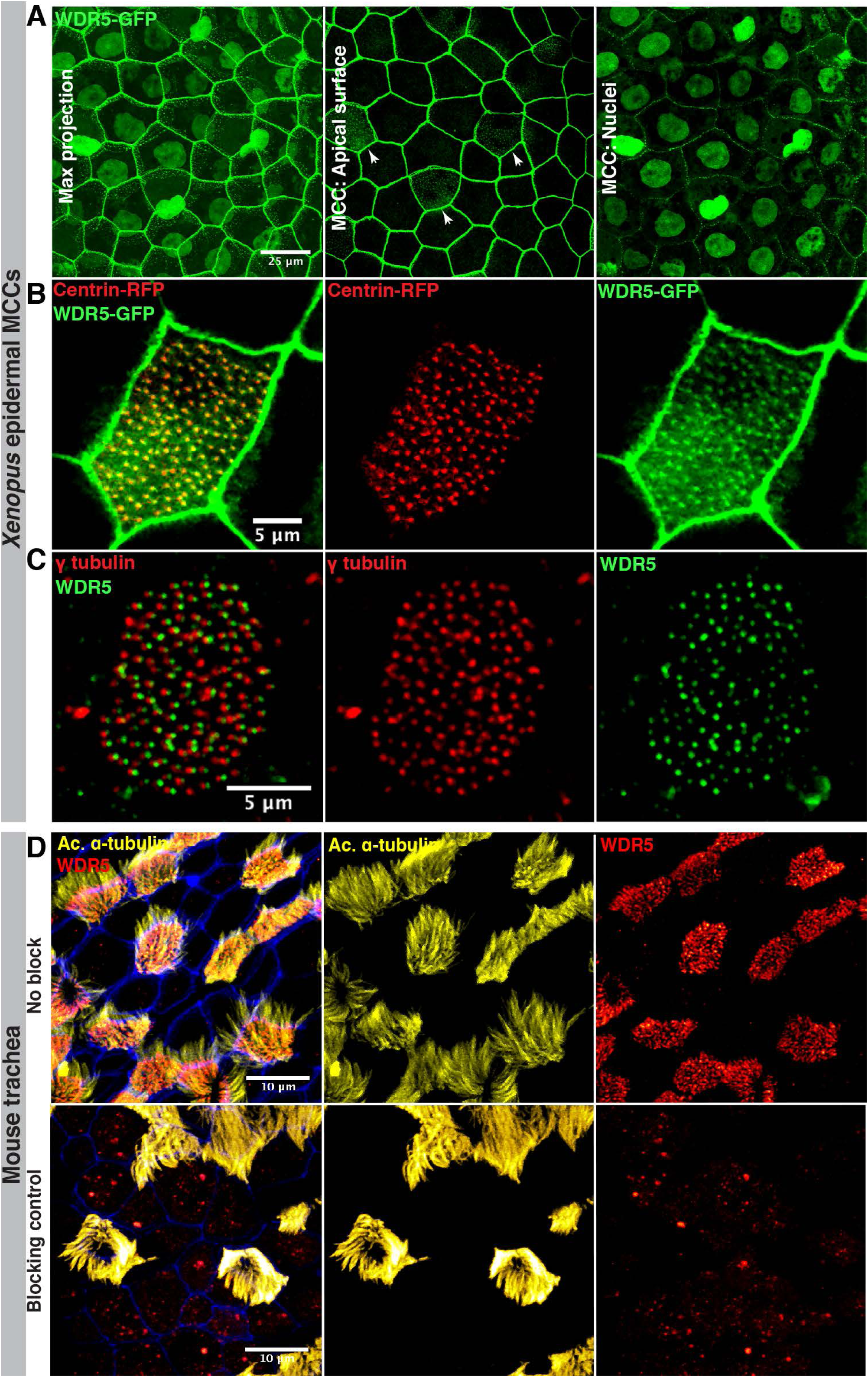
WDR5 is localized near the base of cilia in the MCCs. (A) *X. tropicalis* expressing WDR5-GFP. WDR5 is localized apically in the MCCs in a punctate pattern (white arrows) and in the membrane and nuclei of epithelial cells. (B) *X. tropicalis* epidermal MCC expressing WDR5-GFP (green) and centrin-RFP (red) shows that WDR5 localizes near the basal bodies. (C) Immunofluorescence showing localization of WDR5 (green) and γ-tubulin (red) in a MCC of *X. tropicalis* epidermis. (D) Immunofluorescence showing localization of WDR5 (red), acetylated α-tubulin (yellow) in MCCs of mouse trachea. XY image shows lateral view and XZ image shows orthogonal view. WDR5 signal is lost after incubating the antibody with the blocking peptide. See also Figure S2.

The WDR5 patient mutation, K7Q, fails to rescue cilia driven flow. One possibility is that the K7Q protein variant fails to localize to the base of cilia. To test this hypothesis, we tagged WDR5 (K7Q) with 3xGFP and overexpressed it in *Xenopus* embryos (Figure S2). Contrary to our hypothesis, WDR5 (K7Q)-3xGFP was localized near the bases of cilia suggesting a disruption of WDR5 function rather than localization. Of note, the WDR5 S91K protein variant, which disrupts H3K4MT activity, also localizes near the bases of cilia. Since the WDR5 S91K variant rescues cilia driven flow, we hypothesized that the role of WDR5 in ciliogenesis in MCCs is via its localization at the ciliary base.

### Wdr5 is necessary for basal body patterning and polarization

To understand how WDR5 localization translates into its function, we organized our analysis into four “steps” of ciliogenesis (Figure 3A). The four steps are: 1) MCCs are specified in the deeper epithelial layer, 2) MCCs insert themselves into the superficial epithelia. They simultaneously begin basal body biogenesis within the cytoplasm, 3) Apical enrichment of actin leads to apical expansion of MCCs. Basal bodies also start to migrate and dock to the apical surface, and 4) Basal bodies dock, distribute evenly, and orient at the apical surface, which leads to ciliogenesis. Cilia mediated flow then reinforces basal body polarity. Because WDR5 is localized to basal bodies, we predicted that WDR5 might play an important function in basal body patterning during cilia assembly (step 4). Therefore, we investigated apical docking and the distribution and polarization of basal bodies in *wdr5* morphants.

**Figure 3:**
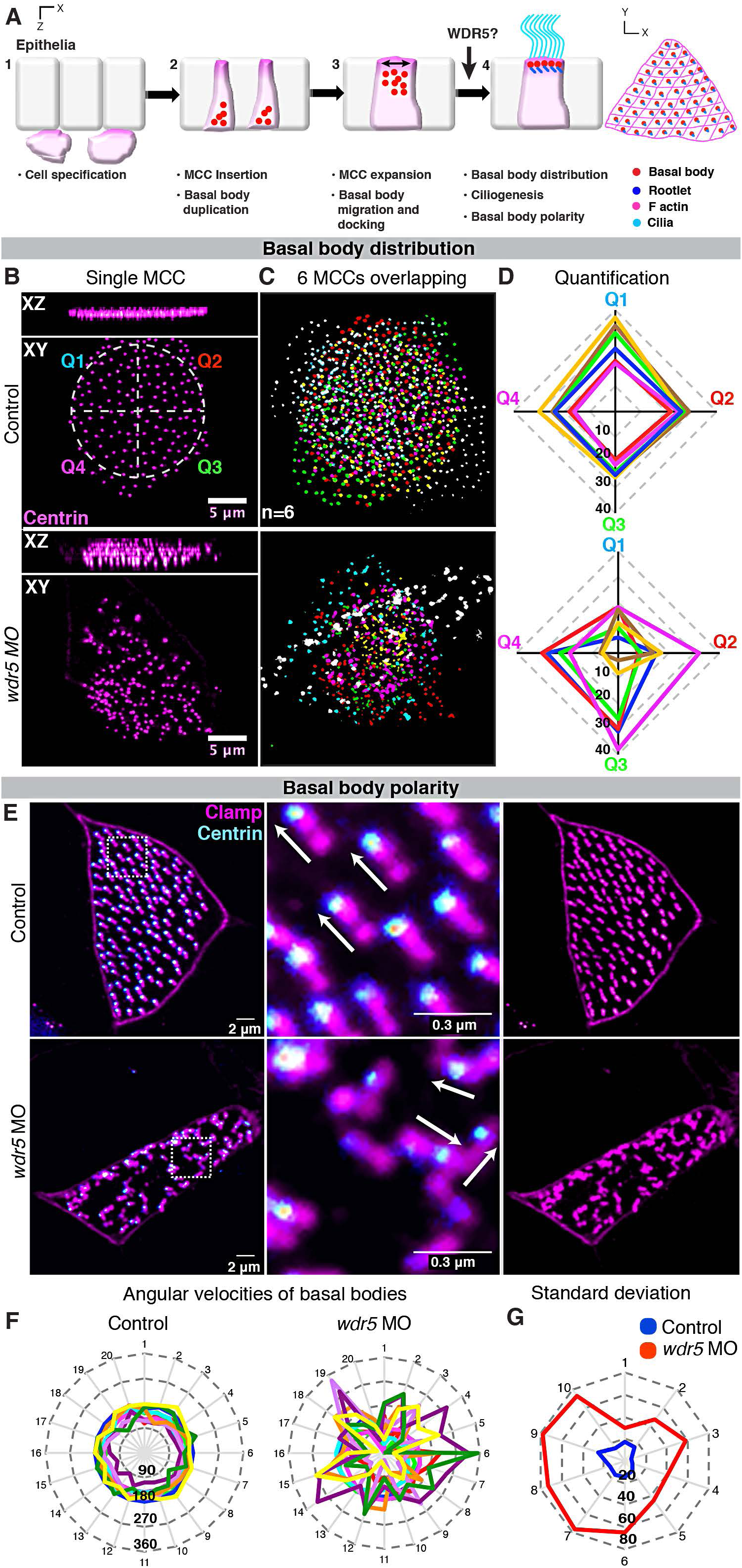
Wdr5 is necessary for uniform distribution and polarization of basal bodies. (A) The four steps of ciliogenesis in MCCs: 1) MCCs are specified in the deeper epithelial layer, 2) MCCs insert themselves into the superficial epithelia. They simultaneously begin basal body biogenesis deep in the cytoplasm, 3) Apical enrichment of actin leads to apical expansion of MCCs. Basal bodies also start to migrate and dock to the apical surface, and 4) Basal bodies dock, distribute evenly, and orient at the apical surface, which leads to ciliogenesis. Cilia mediated flow then reinforces basal body polarity. Wdr5 is essential for uniform basal body distribution in MCCs. (B) MCC showing uniform and clumped distribution of basal bodies (Centrin-RFP) in uninjected control and *wdr5* morphant respectively. Control MCC is divided into 4 quadrants labeled Q1-4 with different colors to use for quantification of distribution pattern in (D). n = number of MCCs. (C) Basal bodies from 6 individual MCCs from different embryos are overlaid on each other to show loss of uniform distribution of basal bodies in *wdr5* morphants. Basal bodies from each MCC are colored differently. (D) Quantitative analysis of basal body distribution in uninjected controls and *wdr5* MO injected embryos. Each MCC was divided into 4 quadrants and the number of basal bodies in each quadrant are plotted on their respective axes (Q1-Q4, see panel D). We measured number of basal bodies in each quadrants for 6 MCCs. In the graph, each color represents one MCC. Squareness represents uniform distribution of basal bodies in a MCC. (E-G) Wdr5 is essential for planar organization of basal bodies in MCCs. (E) MCCs showing parallel and random orientation of rootlets (Clamp-GFP, magenta) with relation to basal bodies (Centrin-RFP – cyan) in controls and *wdr5* MO injected embryos respectively. (F) Quantitative analysis of basal body polarity with angular velocity graphs. Each color represents a MCC (10 total) and axes show orientation of rootlets for 20 basal bodies per MCC. Length of axis represents angle of orientation (0-360). Circularity depicts rootlets are parallel to each other and basal bodies are planar polarized. (G) Angular velocity graph showing standard deviation in the orientation of rootlets. Here, each axis represents each MCC and length represents standard deviation in angle of orientation of 20 rootlets within each MCC. Standard deviation is smaller in controls, as rootlets are more parallel to each other compared to *wdr5* MO injected embryos. ⋆⋆ indicate statistical significance at *P* < 0.005.

We first examined basal body migration and distribution at the apical surface. In WT embryos, centrin-RFP, which marks the basal bodies, is expressed apically and distributed across the cell surface as an ordered array (Figure 3B, upper panels). Depletion of Wdr5 mildly affected apical docking of basal bodies but dramatically disrupted their planar distribution across the apical cell membrane (Figure 3B, lower panels). To visualize the apical basal body distribution defects, we overlaid 6 MCCs (where each color represents a different MCC) from control and *wdr5* morphants (Figure 3C). To quantify their distribution, we divided each MCC into four quadrants and counted the number of basal bodies in each quadrant (Figure 3B,D). In the graph (Figure 3D), each axis represents a different quadrant, the length from the center represents the number of basal bodies in a particular quadrant, and each color identifies a MCC where basal bodies are counted. Control MCCs show similar number of basal bodies in each quadrant (squareness of the plot), whereas *wdr5* morphant MCCs show dramatically different numbers of basal bodies among quadrants (loss of squareness).

Basal body polarity helps drive directional beating and unidirectional fluid flow, which in turn reinforces the polarity of the cilia (Vladar and Axelrod, 2008; Mitchell, et al., 2007). Therefore, in Wdr5 depleted embryos, the loss of cilia and abnormal distribution of basal bodies suggests that the basal bodies may also fail to orient properly. We marked the basal bodies with centrin-RFP and the ciliary rootlets with clamp-GFP to visualize orientation (Figure 3E) (Werner and Mitchell, 2013). In control embryos, the basal bodies and rootlets are organized in an ordered, parallel array (Figure 3E. upper panels). Depletion of Wdr5 disrupts this orientation of the basal bodies and rootlets (Figure 3E, lower panels). We quantified orientation using angular velocity graphs (Figure 3F). The angular velocity measures the orientation of each rootlet, where the vector starts at the end of the rootlet, proceeds along the length of the rootlet, and terminates at the basal body. In Figure 3F, each axis represents the angular velocity of one basal body for a total of 20 basal bodies, where larger angles are plotted further from the center. Each color represents a different MCC for a total of 10 MCCs. In WT cells, the plot is nearly circular, which depicts uniformly polarized basal bodies. The circularity of the plot is clearly disrupted in morphants (Figure 3F). If we measure the standard deviation of the angular velocity in each of the 10 cells, we see that the *wdr5* morphants are dramatically more randomly oriented (Figure 3G). Taken together, we conclude that Wdr5 is necessary for the planar distribution and polarization of the basal bodies, a result that may be related to its location near the ciliary base.

### Wdr5 is essential for apical expansion and actin enrichment in MCCs

To dissect the timing of Wdr5 function, we examined an earlier step (step 3, Figure 4A) of MCC formation. MCCs originate in the basal layers of the epidermis and are then inserted apically to form the superficial epithelia (Stubbs, et al., 2006). Once inserted, the apical surface of the MCC must expand to generate the surface area necessary to host hundreds of cilia (Figure 4A). Measuring apical cell surface area, we discovered that MCCs are inserted but are smaller in *wdr5* morphants suggesting that expansion is affected (Figure 4B,C). Interestingly, the smaller MCC surface area is coupled to a significantly larger surface area in the non-ciliated cells of the embryonic epidermis (Figure 4D), suggesting compensation across the epithelial sheet.

**Figure 4:**
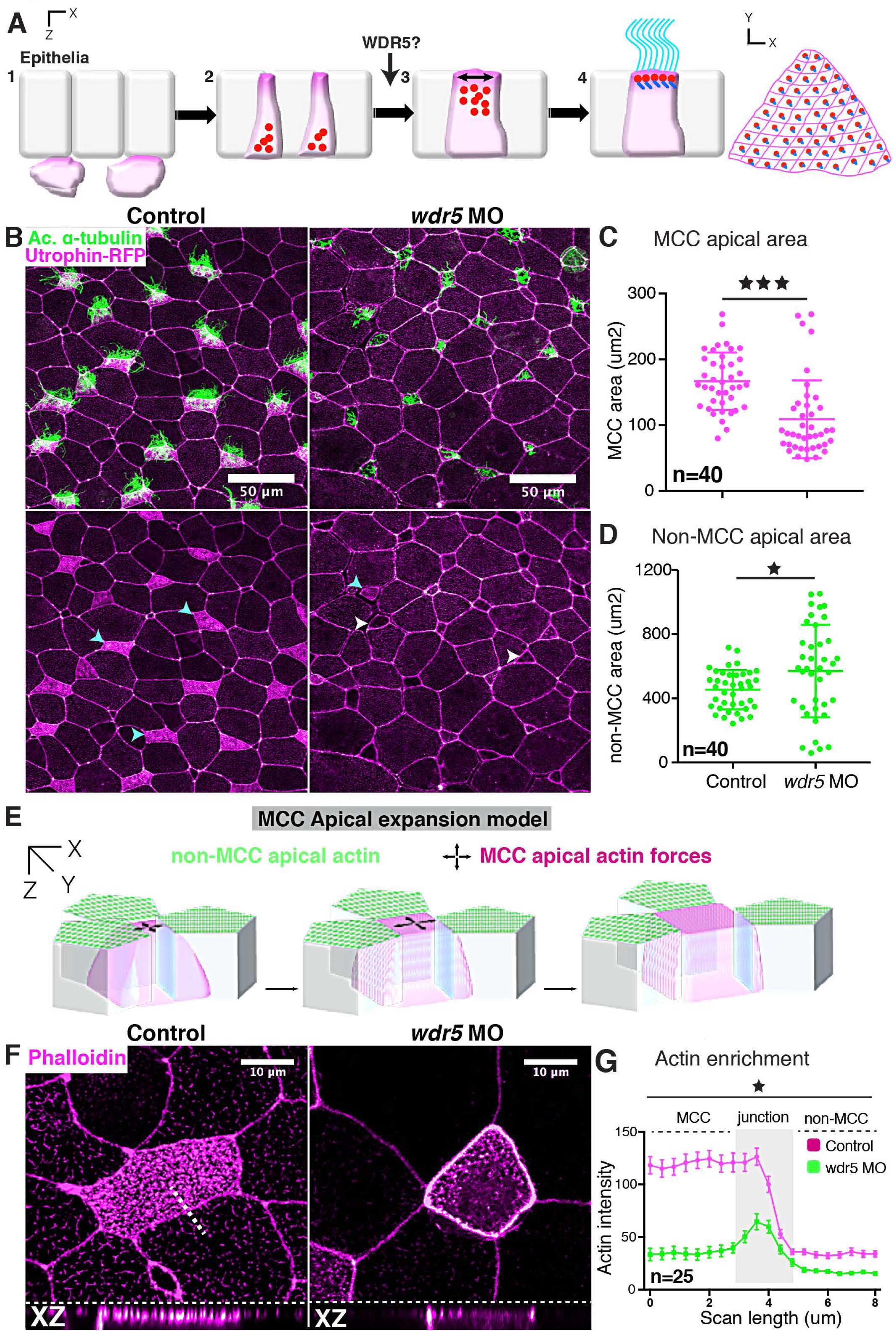
Wdr5 is essential for apical expansion and actin assembly in ciliated cells. (A) The four steps of ciliogenesis in MCCs with emphasis on the role of WDR5 during the third step of MCC formation. (B) *Xenopus* epidermal MCCs marked with anti-acetylated α-tubulin (green, cilia) and utrophinGFP (magenta). Cyan and white arrowheads show ciliated cells with apical enrichment and loss of actin respectively. (C, D) Apical area of MCCs (C) and non-MCCs (D) in controls and *wdr5* MO injected embryos. n = number of MCCs. (E) Model representing actin dependent apical surface expansion in MCCs. (F, G) Quantitative analysis of apical enrichment of actin in MCCs and neighboring non-MCCs in controls and *wdr5* MO injected embryos. Actin enrichment was quantified using a line scan spanning MCC and neighboring non-MCC. Actin was stained using phalloidin. n = number of MCCs. ⋆ and ⋆⋆⋆ indicate statistical significance at *P* < 0.05 and *P* < 0.0005. Data are represented as mean ± SEM.

Little is known about the molecular regulators that are essential for apical expansion of MCCs although the apical actin network plays a prominent role (Sedzinski, et al., 2017; Sedzinski, et al., 2016). Previous work has shown that actin is enriched apically during MCC emergence and is critical for generating cell autonomous 2D pushing forces to expand the MCC (Figure 4E) (Sedzinski, et al., 2016). Given the central role of actin in apical expansion, we wondered if Wdr5 might regulate the enrichment of F-actin at the apical surface of the MCC. We measured F-actin (labeled using phalloidin) in the MCC and the neighboring non-ciliated cells of controls and *wdr5* morphants (Figure 4F,G) using a line scan. In WT embryos, we detected an apical enrichment of actin in the MCCs depicted by high fluorescence intensity; however, loss of Wdr5 led to a significant loss of apical actin (Figure 4F,G). On the other hand, F-actin was only mildly affected in neighboring, non-ciliated cells (Figure 4F,G). Our results suggest that Wdr5 plays a vital role in apical actin enrichment essential for apical expansion and basal body patterning.

**Figure 5:**
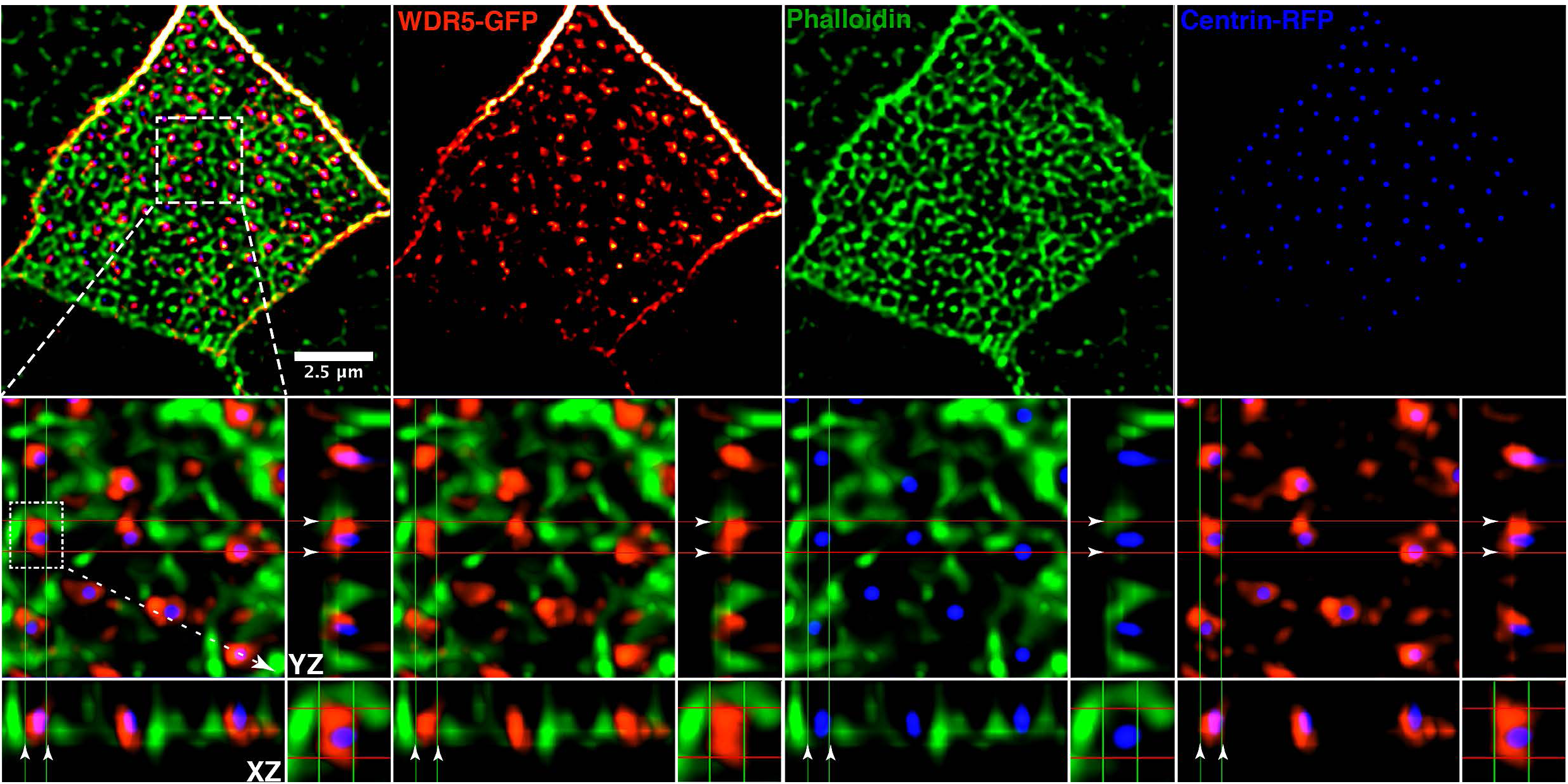
WDR5 is a scaffold connecting basal bodies to actin. A *Xenopus* epidermal MCC expressing WDR5-GFP (red), centrin-RFP (blue) and actin (green) labeled with phalloidin. Basal bodies (blue) dock in the space within the actin lattice. WDR5 localizes between actin and centrin. XZ and YZ projections show orthogonal view. In the orthogonal view, white arrowheads indicate the space between actin and a basal body, which is occupied by WDR5.See also Figure S3.

### WDR5 is interwoven with the actin lattice and connects basal bodies to actin

To connect the non-nuclear localization of Wdr5 (at the ciliary base), basal body patterning defects, and the loss of apical actin in MCCs depleted of Wdr5, we carefully examined the localization of Wdr5, basal bodies, and apical actin. In MCCs, apical actin not only creates the 2D force to expand the apical surface of the cell, it also creates a lattice in which basal bodies are embedded providing a framework for their ordered distribution across the apical cell surface (Sedzinski, et al., 2016; Werner, et al., 2011). Using high-resolution imaging in the MCCs of the *Xenopus* epidermis, we marked human WDR5 with WDR5-GFP, basal bodies with Centrin-RFP, and F-actin with phalloidin (Figure 5). We can readily resolve the apical actin lattice (green) in the MCCs and visualize the basal bodies (blue) docked within the lattice. WDR5-GFP (red) is interwoven within the actin lattice. Interestingly, WDR5 localizes immediately adjacent to basal bodies and the actin network and appears to bridge the gap between the basal bodies and actin. Using optical sections (Figure 5), we can distinguish a clear connection between WDR5 and both the actin filaments and the basal bodies.

To independently investigate a scaffolding function of WDR5 between the basal bodies and the actin lattice, we employed a co-immunoprecipitation assay. We immunoprecipitated endogenous WDR5 from RPE cells and detected both actin and γ-tubulin, which suggests a physical interaction of WDR5 with actin and γ-tubulin (Figure S3).

### The Wdr5-basal body complex interacts dynamically with actin during apical expansion

Given the interaction of WDR5 with F-actin and its importance in organizing apical actin, we hypothesized that WDR5 first localizes apically to organize the actin lattice and then interacts with the basal bodies providing a “docking” site. To test this hypothesis, we used live imaging to monitor these proteins during MCC apical expansion. First we observed WDR5-GFP and to our surprise, WDR5 migrates to the cell surface while apical expansion is proceeding (Figure 6A, Video S2). This process resembled the migration of basal bodies; therefore, we examined localization of basal bodies (Centrin-RFP) and WDR5-GFP concurrently (Figure 6B,C,D). WDR5 was indeed co-localized with basal bodies as they migrated to the apical surface. To gain insight into the relation with F-actin, we visualized F-actin with Utrophin-RFP and concurrently visualized WDR5-GFP (Figure 6E, Video S3, S4). Actin appeared to dynamically organize around WDR5 when WDR5 migrated to the apical surface (Figure 6E, Video S3, S4).

**Figure 6:**
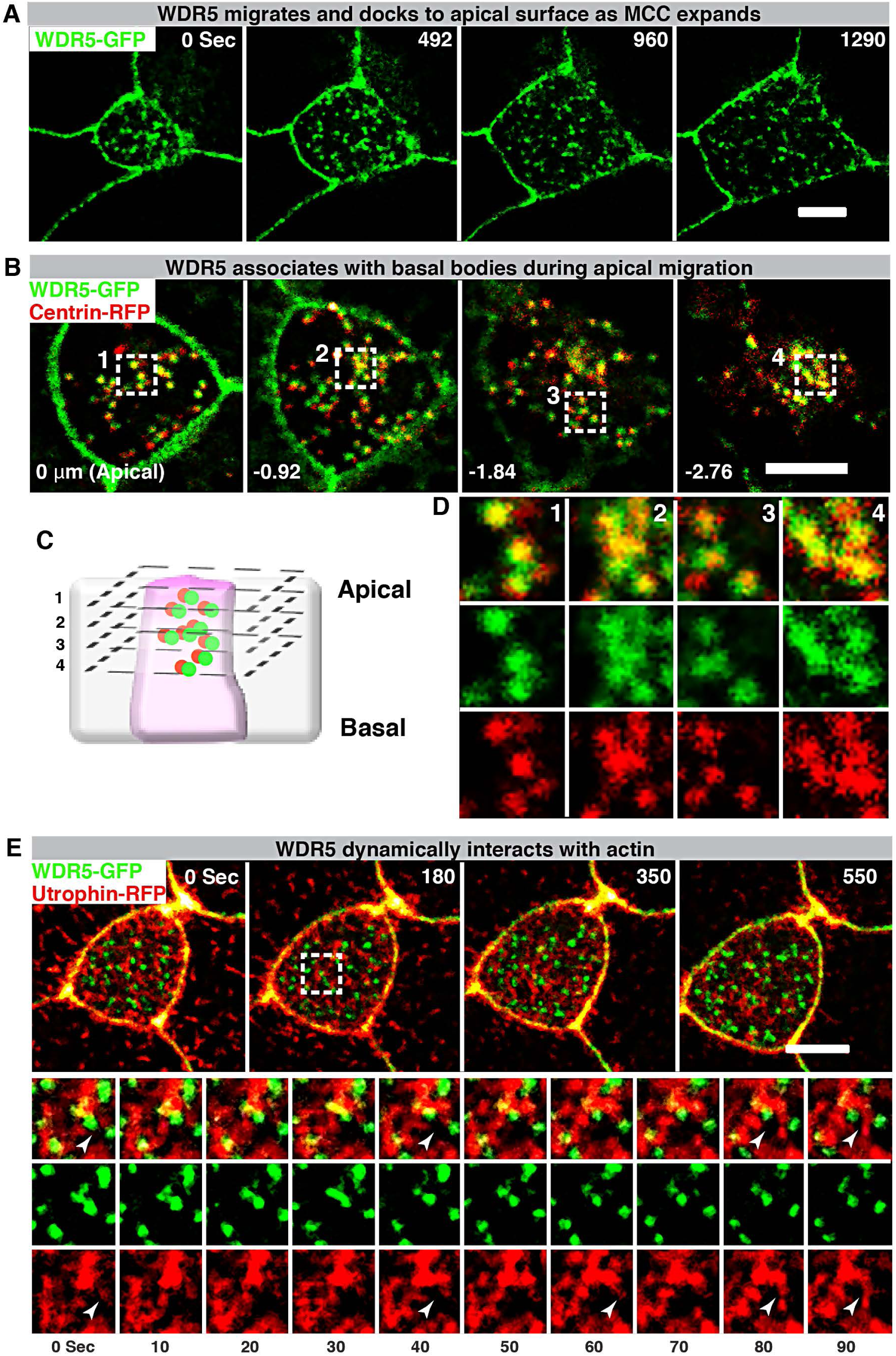
WDR5-basal body complex interacts with F-actin during apical expansion. (A) A montage of *Xenopus* epidermal MCC undergoing apical expansion over 21 minutes. WDR5 marked with WDR5-GFP localizes to the apical membrane as MCC expands. (B,D) A montage of *Xenopus* epidermal MCC expressing WDR5-GFP and centrin-RFP early during MCC expansion process. Basal bodies (centrin-RFP) and WDR5 appear to co-localize from the very beginning when they are deep in the cytoplasm. (C) Schematic showing optical sections of a MCC to examine WDR5 and basal body co-localization in B and D. Optical section 1-4 correspond to the optical sections in the panels B and D. Scale bar = 5μM (E) A montage of *Xenopus* epidermal MCC undergoing apical expansion over 9 minutes. WDR5 (WDR-GFP) interacts with F-actin (Utrophin-RFP) at the apical surface as MCC expands. The region in a white square is zoomed in to show F-actin appears to organize around WDR5. See also Movies S2-S4

### WDR5 stabilizes F-actin in the MCCs

The F-actin network is a dynamic structure and is undergoing continuous assembly and disassembly from G-actin monomers (Blanchoin, et al., 2014). While key proteins like Formins and ARP2/3 are essential for actin assembly, other proteins may play a vital role in regulating the stability of this network by limiting disassembly (Pollard, 2016; Pollard, 1986). Since WDR5 depletion leads to a loss of apical actin enrichment, we hypothesized that either WDR5 promotes the assembly of actin or limits the rate of disassembly. To test these alternatives, we used a Latrunculin A (LatA) based monomer trap for G-actin, which inhibits actin polymerization by binding to G-actin in a stoichiometric 1:1 ratio (Yarmola, et al., 2000; Pollard, 1986). By exposing MCCs to LatA for a specific amount of time and measuring medial actin intensity, we can evaluate the rate of disassembly in MCCs in the presence and absence of Wdr5 (Figure 7A). For this experiment, we injected a suboptimal dose of *wdr5* morpholino (2ng), which only partially affected actin enrichment in the MCCs (Figure 7B). We then exposed the embryos to either DMSO or 2 μm LatA for 10 mins (Figure 7C). We used DMSO treatment as a control for comparisons as actin enrichment in MCCs before and after DMSO treatment was indistinguishable. We compared the normalized medial actin enrichment in MCCs between treatments (Figure 7B). We found that depletion of Wdr5 led to a greater loss of apical actin in response to LatA exposure, consistent with the hypothesis that Wdr5 is essential for stabilizing actin polymers and in turn maintaining the architecture of apical actin in MCCs.

**Figure 7:**
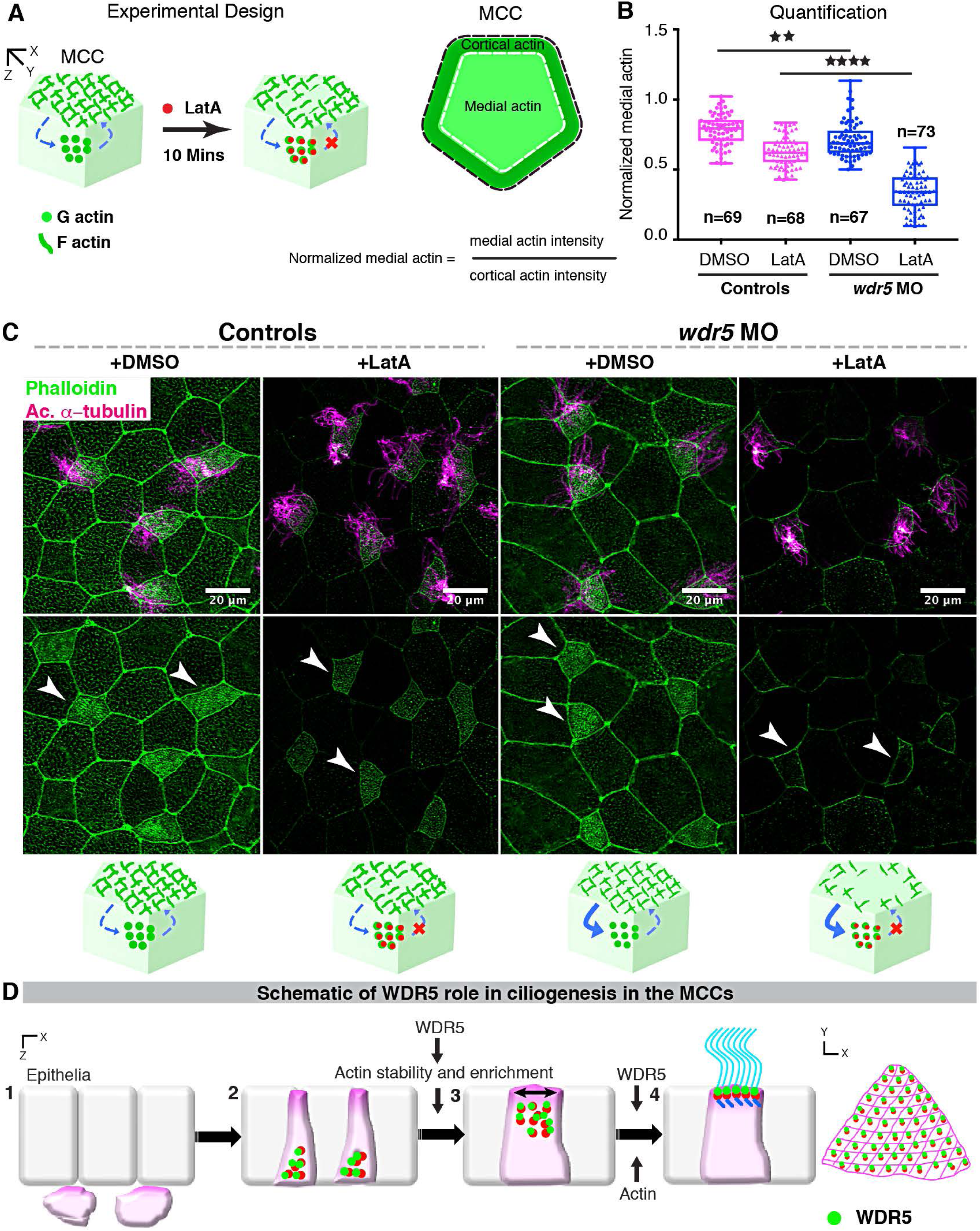
Wdr5 is essential for stabilization of F-actin. (A) Schematic showing the experimental design to examine the rate of F-actin disassembly in MCCs of *Xenopus* embryos. To isolate the effect of WDR5 in disassembly, we exposed epidermal MCCs to Latrunculin A (LatA) treatment for 10 mins. Latrunculin A specifically binds to G-actin in stoichiometric 1:1 ratio preventing the F-actin polymerization. Thus, exposure to LatA prevents the assembly but will not affect disassembly of F-actin allowing us to evaluate the effect of WDR5 depletion on F-actin disassembly in MCCs. Based on earlier results that medial actin is necessary for apical expansion and our results that Wdr5 depletion leads to loss of medial F-actin enrichment, we specifically examined the medial F-actin intensity in controls and *wdr5* morphants. Medial actin intensity was normalized to cortical actin to allow us to combine results of different experiments for statistical comparison. (B) Quantification of normalized medial actin intensity in the MCCs of controls and suboptimal dose of *wdr5* morphants (2nM) after exposure to DMSO (vehicle) or LatA (2uM) for 10 mins. Our results show dramatic reduction in actin intensity in response to LatA in *wdr5* morphants compared to DMSO only. (C) Immunofluorescence showing *Xenopus* epidermal MCCs labeled for F-actin (Phalloidin) and cilia (anti-acetylated α-tubulin) in controls and *wdr5* morphants after exposure to DMSO (vehicle) or LatA (2uM) for 10 mins. Cartoon depicts the hypothesized rates of disassembly and the effect of LatA on medial actin enrichment in controls and *wdr5* morphants. (D) Schematic of a model proposing the role of WDR5 in formation of a MCC. WDR5 binds to basal bodies as basal bodies are synthesized deep in the cytoplasm. WDR5 then migrates apically, where F-actin organizes around WDR5. WDR5 interacts with F-actin to stabilize actin network essential for apical expansion and basal body distribution. In the final stages, WDR5 anchors basal bodies to apical actin network to form functional MCCs.

## DISCUSSION

Our results establish a novel non-chromatin role for WDR5 in F-actin stabilization that is essential for the organization of apical actin in MCCs. Rather than the traditional role for WDR5 as a scaffold for the H3K4 methyltransferase complex, our results suggest an alternative scaffolding function between basal bodies and F-actin in MCCs. We propose the following model for WDR5 in MCC formation and function. First, in MCCs, WDR5 associates with basal bodies and migrates to the apical surface. There it interacts with the developing apical actin lattice to stabilize it, which also generates the force for apical expansion. Stabilization of this actin lattice then provides the organizing framework to uniformly distribute the basal bodies across the apical surface. Finally, WDR5 is essential for anchoring basal bodies to the apical actin network for MCC function (Figure 7D).

Building a MCC is a complex biological process, which involves not only cell migration and intercalation but also dramatic intracellular patterning of organelles for effective function (Meunier and Azimzadeh, 2016; Brooks and Wallingford, 2014). MCCs form in the basal epithelial layer and are inserted into the superficial epithelia. Actin is critical for radial intercalation of cells; however, we did not detect any defects in the apical migration of the MCCs. This suggests specificity in the role of WDR5 in regulating F-actin. WDR5 affects the enrichment of F-actin locally at the apical surface of MCCs, a role that is supported by its localization. In addition, F-actin is hypothesized to bind basal bodies and play a vital role in the apical migration within a MCC (Brooks and Wallingford, 2014; Vladar and Axelrod, 2008; Boisvieux-Ulrich, et al., 1990). While the original study used Cytochalasin D to disrupt actin and showed that basal bodies arrest in the cytoplasm, they did not provide any direct evidence like co-localization of F-actin and basal bodies in the cytoplasm (Boisvieux-Ulrich, et al., 1990). We did not observe any F-actin bound to apically migrating basal bodies. Further, Wdr5 depletion led to relatively small defects in basal body apical migration. As Wdr5 depletion is partial, whether the remaining actin is sufficient or whether actin does not have a role in basal body migration requires further testing. Clearly, WDR5 and actin are essential for the apical expansion, planar distribution and polarity of the basal bodies, and ciliogenesis.

F-actin organizes into a network that is essential for apical expansion (Sedzinski, et al., 2016; Stubbs, et al., 2006). Given that F-actin is inherently dynamic, the molecules that stabilize F-actin are necessary to maintain the actin lattice in the MCCs. Using live imaging and an actin monomer trap, we discovered that WDR5 is essential to stabilize F-actin in MCCs. Importantly, Wdr5 first associates with basal bodies before they dock to the apical surface. Then, Wdr5 interacts with actin polymers to stabilize them. The inter-relationship between apical actin assembly and basal body migration and docking is still unclear, however, numerous studies have found that basal body migration/docking defects are often associated with failure to enrich apical actin (Epting, et al., 2015; Antoniades, et al., 2014; Brooks and Wallingford, 2014; Ioannou, et al., 2013; Park, et al., 2008). Our results suggest that the process of basal body migration recruits molecules such as WDR5 to stabilize the apical actin network. Interestingly, molecules essential for actin remodeling are often localized to the basal bodies (Epting, et al., 2015; Antoniades, et al., 2014; Park, et al., 2008; Huang, et al., 2003) suggesting apical actin lattice organization is coordinated by apical migration of basal bodies.

Finally, our results emphasize the importance of patient driven mechanism discovery. Recent studies in CHD and autism clearly point to a role for chromatin modifiers in disease pathogenesis, but the molecular mechanisms remain unclear (De Rubeis, et al., 2014; Zaidi, et al., 2013). WDR5 was identified as a candidate CHD/Htx gene; given the respiratory complications associated with these diseases, we sought to examine MCC function in our high-throughput *Xenopus* mucociliary disease model(Garrod, et al., 2014; Werner and Mitchell, 2013; Nakhleh, et al., 2012; Swisher, et al., 2011; Tan, et al., 2007; Houtmeyers, et al., 1999). We identified an unexpected non-chromatin function for WDR5 in mucociliary clearance providing a plausible pathogenesis mechanism for respiratory complications in patients with WDR5 related CHD/Htx. Thus, the discovery of a cilia phenotype may have important clinical implications for patient management.

## ACKNOWLEDGEMENTS

We thank the patients and their families who are the inspiration for this study. We thank Sarah Kubek and Michael Slocum for animal husbandry. Thanks to the Center for Cellular and Molecular Imaging at Yale for confocal imaging. The authors thank Dr. Ann Miller and Dr. Michael Cosgrove for the C-3XGFP and human WT and S91K- WDR5 constructs respectively. The authors with to thank Prof. Thomas Pollard for helpful discussions on actin biology and Shiaulou Yuan and Emily Legue for helpful comments on the manuscript. This work is supported by an NIH/NHLBI K99/R00 award to SSK. This work was supported by a Pilot Project as part of NIH/NHLBI 5U01HL098188 and NIH/NICHD R01HD081379 and NIH/NHLBI R33HL120783 to MKK. MKK is a Mallinckrodt Scholar. The contents of this publication are solely the responsibility of the authors and do not necessarily represent the official views of the NHLBI. Please see the supplementary material for additional acknowledgements.

## AUTHOR CONTRIBUTIONS

SSK and MKK conceived the work, designed the experiments, interpreted all experimental data, and wrote the manuscript. SSK performed all the experiments except JNG performed Western blots to determine WDR5 levels and interpreted this data. KFL dissected mouse tracheas for immunofluorescence. All authors reviewed and contributed to the writing of the manuscript.

## METHODS

### Animal husbandry

*Xenopus tropicalis* were housed and cared for in our aquatics facility according to established protocols that were approved by the Yale Institutional Animal Care and Use Committee (IACUC). Mice were also housed and cared for according to established protocols that were approved by the Yale IACUC.

### Microinjection of morpholino oligonucleotides and mRNA in *Xenopus*

Embryos were produced by *in vitro* fertilization and raised to appropriate stages in 1/9MR + gentamycin according to established protocols (del Viso and Khokha, 2012; Khokha, et al., 2002). Staging of *Xenopus* tadpoles was as previously described (Nieuwkoop, 1994). Morpholino oligonucleotides or mRNA were injected into one-cell or two-cell embryos as described previously (Khokha, et al., 2002). The following morpholino oligonucleotide was injected: *wdr5* translation blocking (2.5-4 ng/embryo 5’-CGGGTTTCTTTTCTTCTGTTGCCAT-3’). The scrambled MO (4ng/embryo 5’-CCTCTTACCTCAGTTACAATTTATA -3’) was injected as a negative control. Alexa488 (Invitrogen), mini-ruby (Invitrogen) or green fluorescent protein (100 pg/embryo) were injected as tracers. We generated *in vitro* capped mRNA using the mMessage machine kit (Ambion) and followed the manufacturer’s instructions. Full length human WDR5 was purchased from Thermo Scientific (IMAGE clone: 3538255) and subcloned into the pCSDest vector using Gateway recombination techniques. S91K-WDR5 was a kind gift from Dr. Cosgrove, SUNY Upstate Medical University. We generated the K7Q mutation using PCR amplification. C-3xGFP was a gift from Dr. Ann Miller, University of Michigan, Ann Arbor. Human WT-WDR5, S91K-WDR5 and K7Q-WDR5 were PCR cloned in frame into the 3xGFP vector. We injected 400pg wild type human WT-WDR5, WDR5-GFP, K7Q-3xGFP and S91KWDR5 RNA for rescue of *wdr5* morphants. Centrin-RFP (100pg), Clamp-GFP (150pg), memRFP/GFP (150pg), WDR5-GFP (200pg), and utrophin-RFP/GFP (150pg) were injected into one-cell embryos to mark basal bodies, ciliary rootlets, cilia and membrane, WDR5, and F-actin respectively. Latrunculin A was a gift from Dr. Thomas Pollard at Yale University. Latrunculin A was dissolved in DMSO and used at a final concentration of 2 μM.

### Immunofluorescence

We harvested mouse trachea from euthanized adult mice. Mouse tracheas were fixed in 4% paraformaldehyde/PBS overnight at 4^°^C when we assayed for acetylated α-tubulin and WDR5. For immunofluorescence of multiciliated epidermal cells, we used stage 28–30 *Xenopus* embryos. *Xenopus* embryos were fixed in 100% chilled methanol overnight at -20^°^C for γ-tubulin and WDR5. Mouse trachea and *Xenopus* embryos were mounted in Pro-Long Gold (Invitrogen) before imaging. WDR5 blocking peptide (Bethyl) was used to examine WDR5 specificity in 1:1 and 1:2 (antibody: blocking peptide) ratio.

### Image analysis

Images were captured using a Zeiss 710 Live or Leica SP8 confocal microscope. Images were processed in Fiji, Image J, or Adobe Photoshop. Figure 5 was deconvolved using Huygens Professional to increase the clarity of the signal. The 3D and orthogonal projections of Fig 5 was generated using Leica LAS X software. Basal body polarity was measured using Fiji. Final figures were made in Adobe illustrator.

### Statistical analysis

Statistical analysis was done using PRISM, JMP and Vassarstats software. All the comparisons were made using t-tests after confirming the normal distribution of the data. We randomly picked one cell *X. tropicalis* embryos from fertilization as uninjected controls or for morpholino or RNA injections. Investigators were not blind to experiments or the statistical analysis.

### Antibodies

Mouse monoclonal Anti-acetylated α-tubulin (Sigma, T-6793; 1:1000), Rabbit polyclonal Anti-WDR5 (Bethyl, A302-430A; 1:150 for immunofluorescence, 1:400-1:500 for immunoprecipitation and 1:500-1:1000 for western blot), Rabbit polyclonal Anti-WDR5 (Bethyl A302-429A; 1:400-1:500 for immunoprecipitation), Goat polyclonal Anti-Actin (Santacruz, SC-1615; 1:250 for western blot), Mouse monoclonal Anti-γ tubulin (Sigma, T6557; 1:100 for immunofluorescence, and 1:4000 for western blot), Mouse monoclonal Anti-GAPDH (Ambion, AM4300; 1:10,000 for western blot), HRP tagged secondary antibodies (Jackson Immuno Research Laboratories, rabbit 1:12000, mouse 1:16000, goat 1:5000: for western blot) were used. Alexa 488, Texas red and Alexa-647 (all 1:500) were used as secondary antibodies for immunofluorescence. Alexa 633 and 488 phalloidin (both 1:40) were used. Blocking peptide for WDR5 was purchased from Bethyl for antibody A302-430A.

### Co-immunoprecipitation

For immunoprecipitation, RPE cells were grown until 100% confluence and then were starved for 24-48 hours before lysis. Cells were lysed with NP-40 buffer (150 mM NaCl, 1.0% NP-40, 50 mM Tris, pH 8.0) with protease inhibitor on ice at a concentration of 1ml of buffer for a 10 cm plate. Cells were then centrifuged to collect supernatant. Supernatant was then pre-incubated with protein A/G beads for 1-3 hours to eliminate unspecific binding. The mixture with beads was centrifuged to collect supernatant. Supernatant was incubated with a respective antibody for 1-2 hrs, followed by overnight incubation with beads at 4^°^C. Lysate was then centrifuged the following day and the supernatant was discarded, and the beads were collected and washed with 0.1% PBS-Tween. 2x SDS loading dye was then added to the beads and the samples were analyzed with Western blot. Western Blot was carried out using standard protocol with 4-12% gels.

### Protein extraction for Western Blot analysis

To obtain total cell lysate, pools of 20 staged controls, *wdr5* morphants or *wdr5* MO + human WT WDR5 RNA injected *Xenopus* embryos were placed in 200 μl of 1x RIPA buffer and crushed using a pestle. Samples were spun down twice (10,000g rpm for 20 and 10 mins at 4o degrees) to remove fat and debris. Protein samples were used immediately for western blot or stored at -80^°^ degrees). Western Blot was carried out using standard protocol with 4-12% gels. Quantifications of changes in protein level were calculated using ImageJ software from NIH.

